# Feasibility of a parent-delivered attention and working memory intervention for early school-aged children born preterm

**DOI:** 10.1101/2025.10.10.680956

**Authors:** Signe Bray, Tao Tao, Sakshi Kaur, Mervyn Singh, Amanda Ip, Shelly Yin, Daria Merrikh, Stella Heo, Ella Ryan, Sunny Guo, Leonora Hendson, Deborah Dewey, Sarah Macoun

## Abstract

Preterm birth is a common neurodevelopmental condition that can have lasting impacts on cognition, including attention and working memory. Interventions that strengthen these skills in early childhood could support school readiness. Dino Island is a tablet-based intervention that combines process-specific practice of attention and working memory skills with a compensatory component that teaches metacognitive skills to scaffold learning. In this pilot study, we evaluated the feasibility of parent-delivered Dino Island in young children born preterm (N=10), alongside a control group who played educational games (N=12). Both groups were instructed to play 2-3 times per week for approximately 20 minutes over a 12-week period. Attention and working memory on untrained tasks were assessed before and after completing the intervention. Parents provided fidelity data through tracking sheets and participated in exit interviews to offer feedback and identify barriers and facilitators. We found that the Dino Island program was successfully delivered by parents with high fidelity. Attrition was higher in the Dino Island group, likely reflecting the challenges of delivering cognitive remediation in home settings. Comparison of attention and working memory scores on untrained tasks pre- and post-showed practice effects but no specific benefit of Dino Island. However, parent reports suggested behavioral improvements specific to the Dino Island group, noting far-transfer effects where children applied metacognitive strategies in other contexts. Overall, this work shows feasibility and tolerability of Dino Island in young children born preterm. Future research should examine its potential impact on school readiness and longer-term academic outcomes in this population.

## Introduction

Preterm birth occurs in an estimated 10% of live births (Ohuma et al., 2023) and is associated with elevated risks for long-term neurodevelopmental challenges, including persistent difficulties with attention, executive functioning, and learning (C. S. Aarnoudse-Moens et al., 2009; Fraiman et al., 2023; McBryde et al., 2020; Robinson et al., 2023; Sucksdorff et al., 2015). Children born extremely or very preterm (< 32 weeks gestational age) are two to three times more likely to receive a diagnosis of attention-deficit/hyperactivity disorder (ADHD) compared to their term-born peers (Bhutta et al., 2002). While children born extremely or very preterm are often the focus of developmental follow-up, emerging evidence indicates that those born moderate-to-late preterm (33–36 weeks gestational age) also exhibit subtle but meaningful cognitive and behavioral vulnerabilities (Martínez-Nadal & Bosch, 2020; Mitha et al., 2024; Woythaler, 2019). These difficulties, particularly in attention regulation and executive control, contribute to lower levels of school readiness among preterm-born children and increase their risk for academic underachievement and broader educational challenges upon school entry (Karnati et al., 2020; Lock et al., 2024).

Interventions aimed at strengthening attention and executive function during early childhood may be particularly impactful in promoting school readiness in this population. There is evidence from several studies and samples suggesting that cognitive training programmes can effectively improve attention and executive function and have the greatest benefits during early developmental windows (Karbach & Kray, 2009; Rueda et al., 2012; Scionti et al., 2019; Wass, 2015). In particular, digital educational games have been shown to significantly improve attention and related cognitive skills in typically developing preschool-aged children, highlighting the potential of ‘serious-games’ as an engaging and accessible modality for early intervention (Prins et al., 2011; Sop & Hançer, 2024).

With this in mind, Dino Island (DI) was developed as a tablet-delivered cognitive intervention designed to support building attention and executive functioning skills in young children. This program is based on an earlier version of the program called Caribbean Quest (CQ) which has shown preliminary efficacy in strengthening cognitive abilities in children with neurodevelopmental diagnoses (Macoun et al., 2022; Macoun et al., 2021). The DI intervention is a hybrid intervention based combining process specific (Mateer et al., 1996; Sohlberg et al., 2003) and compensatory (Cicerone et al., 2011) approaches which may be more effective than either approach alone with respect to outcomes and generalization (Cicerone et al., 2011; Kennedy et al., 2008; Partanen et al., 2015; Schwaighofer et al., 2015; von Bastian & Oberauer, 2014). Games for younger children within DI target sustained attention and working memory, and are designed to be self-adjusting and hierarchical, providing massed practice and intense exercise to maximize plasticity (Kleim & Jones, 2008). The intervention is delivered by a trained non-expert interventionist (e.g., any adult in the child’s circle of care) who supports their child in learning metacognitive strategies such as good listening and self-awareness. While this program has shown promise in several neurodevelopmental populations (Kerns et al., 2010; Macoun et al., 2022; Macoun et al., 2021), no studies to our knowledge have tested the efficacy of a combined process-specific and compensatory approach in supporting attention and working memory in young children who were born preterm.

The DI/CQ program must be delivered by an interventionist who is trained on supporting children’s use of metacognitive strategies. Parent-delivered interventions can provide secondary benefits such as enhanced generalization (Mingebach et al., 2018) and enhanced parent-child communication (Brian et al., 2022). Research also demonstrates that when parents are equipped with evidence-based strategies and are supported in applying them consistently at home, they can support their child’s development in meaningful ways (Kaminski et al., 2008). In this study, parents were trained using a standardized script and fidelity was monitored via video calls during which application of meta-cognitive strategies was assessed and feedback provided. Outcomes of interest in this work were whether parents could successfully implement metacognitive strategies when working with their children and whether upon completion, they perceived benefits for their children.

The goals of DI are to support attention and working memory skills beyond the games practiced as part of the intervention. That is, the intervention is designed to support both near- and far-transfer effects. Here we chose a battery of attention and working memory tests administered before and shortly after completion of the intervention. In addition, parents completed an exit interview in order to assess potential changes in behavior that are more difficult to capture in a lab-based setting. A challenge with cognitive intervention research is that participants can show improvements related to practice and increased familiarity with the tasks used to assess transfer (Ripp et al., 2022). Moreover, the interaction with interventionists may also support behavior changes that are non-specific to the intervention itself. We therefore chose to use an educational games control group in this study that would allow us to control for practice effects and exposure to a parent-led tablet-based game activity. Games for the control group were chosen to be educational, e.g. focused on early literacy or numeracy skills, but to not include any challenge to attention and working memory. The Educational Games (EG) group did not receive coaching on metacognitive strategies.

In the work reported here, we tested the feasibility, acceptability and efficacy of a parent-delivered DI intervention in a small sample of 4- to 6-year-old children who were born prematurely. We compared outcomes from DI to an EG control group. Our findings can contribute to the ongoing refinement of programs such as DI and support future intervention work targeting cognitive skills in children born preterm.

## Methods

### Recruitment

Children aged 4-6 years and their parents were recruited in Calgary, Alberta, and surrounding areas to participate in a study on brain and cognitive development in children born prematurely, involving MRIs, cognitive testing and questionnaire data. Families recruited into the larger study were offered the opportunity to participate in the intervention sub-study described here. Participants were recruited through the Alberta Children’s Hospital Neonatal Follow-Up Clinic and community advertisements including social media. Recruitment occurred between December 2018-May 2022. To meet inclusion criteria, children must have been born preterm (PT; < 37 weeks completed gestational age), be fluent in English, and have at least one parent or caregiver who was also fluent in English and able to deliver the active or control intervention. Children were excluded if they had MRI contraindications, a Full-Scale Intelligence Quotient (FSIQ) < 70, bilateral blindness with corrected visual acuity <20/200 in the better eye, seizure disorder, bilateral neurosensory hearing loss, cerebral palsy or history of severe head trauma. Behavioral diagnoses such as autism or ADHD were not exclusionary.

A total of 72 families recruited into the larger study were offered the opportunity to participate in this sub-study, and of those, N=35 consented to participate. The first three families were assigned to the DI group, for the study team to become familiarized with training parents on meta-cognitive strategies. Following that, families who agreed to participate in the intervention were assigned to the DI or EG groups using a randomized assignment generated using graphpad.com. This resulted in N=20 children assigned to DI, and N=15 children assigned to the educational games control group.

Thirteen children withdrew before completing the study, nine voluntarily (N=8 in the DI and N=1 in the EG) and four (N=2 DI and N=2 EG) due to the COVID-19 pandemic, which resulted in insufficient intervention hours, or missing data on pre- and/or post-testing. As a result, the final sample for analysis consisted of 22 children, comprising 10 in the DI and 12 in the EG groups.

Study procedures were approved by the University of Calgary Conjoint Health Research Ethics Board (CHREB) #REB18-0876. Parents provided informed consent and children provided assent on REB approved forms.

## Study Design

This study used a single-blind design wherein the research team had knowledge of group assignment while parents and children were not informed which group they were assigned to. Parents were told that in this project we were testing two types of educational games, they would be randomly assigned to one type of game, and researchers did not know if one type of game offered more benefits for children. Children completed a baseline neuropsychological assessment (∼2.5 hours) to obtain pre-intervention cognitive and behavioural data. Neuropsychological tests were administered by trained research assistants who were not blind to group membership.

The intervention was parent-delivered and training for parents was done through a PowerPoint presentation delivered by the research team in-person with a question-and-answer period, with content specific to each type of intervention. Both interventions were tablet-based, and families were provided with tablets for the duration of the study. Parents were instructed to deliver the intervention at home over a period of 12 weeks (2-3x a week for 20-30 minutes session) for a total of 12 recommended intervention hours, divided equally across different games. Parents were instructed to remain with their child 100% of the time during game play. In the DI group, parents were instructed to work with their children to deliver metacognitive strategies. Duration and intensity were based on similar research in child populations (Kerns et al., 2010; Macoun et al., 2022; Macoun et al., 2021). At least once during the intervention, a research assistant completed a video observation with families and assessed fidelity as well as provided feedback to families on administration of the interventions. Following the 12-weeks of intervention, children returned for post-testing with the same neuropsychological battery. Our design included a control group to mitigate the impact of potential practice effects on findings.

### Dino Island Intervention

DI is the latest iteration of a tablet-based serious game designed specifically for children (Kerns et al., 2017; Kerns et al., 2010; Macoun et al., 2022; Macoun et al., 2021). It combines process-specific attention/executive function training in five therapeutic games designed to strengthen sustained attention, selective attention, visual/verbal working memory, and attention switching/flexibility. DI is delivered by a trained interventionist (here, a parent or caregiver) who supports the child in applying metacognitive strategies during game play. Parent training included instruction in the following areas: 1) education about attention and executive function, 2) education about the approach used with the DI intervention, 3) training on positive behavioural strategies, 4) training on how to teach metacognitive strategies, 5) and training on how to download and play the DI games. Only three of the five games (focused on sustained attention and working memory) were deemed suitable for preschool-aged children and were recommended to play in the current study: Mining Cave, Wave and Shell Machine.

*Mining Cave* specifically targets sustained attention. This game starts by showing the child one or more dinosaurs they must remember. The specific rules and the dinosaurs that need to be remembered/selected change across levels, so children must carefully follow each level’s instructions.

Children see a long line of dinosaurs walking by and their job is to hit the big red button on the screen anytime the target dinosaur(s) walk underneath the light on the cave ceiling. On more difficult levels, the child is faced with distractions (sounds and creatures crawling around the cave), more complex instruction sets, and part of the screen is visually blocked by rock pillars, so they must retain information longer and have less time to respond. Later levels of mining cave involve n-1 and n-2 back tasks where children must press the button if the dinosaur is the same as the one that came either 1 or 2 before. Children must complete the task to 90% accuracy in order to progress to the next difficulty level.

*Wave* focuses on strengthening impulse control and sustained attention, particularly at early levels. During the game, children ride a pterodactyl over the ocean and must collect specific items, while avoiding nontarget items. The pterodactyl must be positioned directly over the object to collect each item. The child must steer their pterodactyl away from incorrect items. As the game progresses, levels become longer and with more distractions, more items to remember and more complex instruction sets, making the game increasingly more difficult. Children must complete the game to 90% accuracy to progress to the next difficulty level.

*Shell Machine* exercises attention and visuospatial/auditory working memory. On Shell Machine, the child is instructed to drag shapes, held by dinosaurs, to a shell machine in the order that they inflate. The game begins with simple visual forward span tasks where children are required to remember a series of items in order. It progresses to visual backward span tasks where they must remember a series of items in reverse order and subsequently to auditory forward and backward span tasks. The span length increases in difficulty when a child obtains three correct items in a row and decreases in difficulty when they make one error. In later levels, the dinosaurs/items change position on the screen. Children must pass five spans in order to progress to the next difficulty level. Difficulty levels increase by increasing span length and increasing the complexity of the task (e.g., reverse spans, spans where they must ignore certain types of items and moving items).

All games are self-adjusting and hierarchical, providing massed practice and intense exercise. Game progression is dependent upon either achieving a specific number of correct trials (100% correct on five span lengths as in the Shell Machine game) or a certain accuracy level (90% accuracy level as in Mining cave), depending upon the specific game and cognitive process being targeted. Between the attention and working memory games, children are rewarded with fun bonus games to enhance motivation, where they are able to collect coins and buy costumes to decorate their avatar.

#### Metacognitive strategies

A unique element of the DI program is the teaching and use of metacognitive strategies. Parents were taught how to use various metacognitive strategies including: self-awareness, good listening, self-control, moving your body, rehearsal, elaboration, chunking, visualization, substituting/paraphrasing, and tracing. These strategies focus on engaging in problem solving and self-regulation, and were taught as part of the DI program and delivered by the parental interventionist during the child’s game play. Additionally, parents were encouraged to use each session as an opportunity to help their child improve in areas like resilience by completing each game to the best of their ability while also developing a sense of time by indicating game days on a calendar throughout the study.

### Educational Games Control Condition

Children in the control group played age-appropriate educational games throughout the 12-week intervention period for the same duration as the intervention group (20 minutes a day, 2-3 times a week). Games were chosen to allow one-on-one interaction between parents and their child, mirroring the DI intervention, but not to directly enhance attention or working memory abilities. Specific games included Elmo loves 123, ABC Mouse, Khan kids, Puzzle kids, and 123 Numbers, Colours, and Shapes. This condition also controlled for effects of typical attention development, repeat assessments, and playing a tablet-based game under parent supervision. Parent training in this condition focused on building a consistent schedule for gameplay, keeping their child motivated by immediately providing rewards and creating an optimal and controlled environment that was tidy, quiet, and comfortable for the child. As in the DI group, parents were encouraged to use each session as an opportunity to support resilience and work with their children to track days played on a calendar. The training session also instructed parents on how to fill out the tracking sheet, as well as information about future surveys and calls to check in on the child’s progress and ensure that the sessions were conducted in a controlled manner.

### Tracking progress and exit interviews

For both the intervention and control groups, parents completed tracking forms and an exit interview to assess fidelity and gather feedback to support program development.

#### Tracking forms

Parents were asked to complete tracking forms during each session, which provided data about which games were played, how long in each session, what levels were achieved and in the DI condition, information about metacognitive strategies applied during the sessions.

To analyze these data, for each participant we counted the total number of tracking sheets and total time spent on the games. These values were compared between the DI and EG groups using unpaired t-tests to determine whether groups received approximately equal amounts of intervention.

For the DI group only, metacognitive strategy use was tallied from the tracking forms. We report the total number of strategies reported per family along with the average number each family reported using per session. To describe which strategies were used most frequently, we divided the total number of reports for each strategy (within and across individuals) by the total number of strategies reported.

#### Exit interviews

After completing the intervention, an exit interview form was given with families to provide feedback and comments about their experience with the intervention. This questionnaire asked parents whether they noticed any changes in their child’s behaviour in various areas like attention, social skills, and remembering. Furthermore, the form asked parents to share whether their child was using any metacognitive strategies. We also asked about their experiences in delivering the programs and any barriers/facilitators they experienced. We compared the frequency with which specific behavior changes were reported in each group, and summarized parent feedback from the two groups.

### Neuropsychological Measures

Data was collected across several cognitive and behavioural domains at baseline and immediately following the intervention.

#### Global intellectual functioning

Intellectual ability was assessed using the Wechsler Preschool and Primary Scale of Intelligence (WPPSI-IV-CDN) (Wechsler, 2012). The WPPSI-IV-CDN is a commonly used standardized measure that provides an overall estimate of full-scale intellectual ability (Wechsler, 2002). Raw scores from this measure were transformed into standard scores with a mean score of 100 and a standard deviation score of 15. The WPPSI was only administered at the baseline visit; scores are reported for descriptive purposes to characterize the sample.

#### Behavior challenges

The Conner’s Early Childhood (Conners, 2009) was completed by parents at baseline and used to characterize behavior challenges as well as assess any differences between the DI and EG groups prior to starting the intervention.

#### Visual Acuity

To assess visual acuity, all children were screened using the Goodlite Crowding Cards (Good-Lite). This measure was administered given the high risk of visual deficits in PT children to ensure that children had sufficient visual acuity to participate in the DI and control conditions. All children passed the visual acuity assessment.

### Attention and Executive Functioning Measures Administered Pre- and Post-Intervention

Children completed a number of assessments at baseline and post-intervention follow-up.

#### Cognitive Flexibility

The Dimensional Change Card Sort Task (DCCS) is a commonly used measure for assessing executive function early in development (Zelazo, 2006). The DCCS has strong validity and reliability and is appropriate for the target age-range in the current study (Zelazo, 2006). The task involves learning to sort a series of laminated cards into containers first by color (6 trials), next by shape (6 trials), then a more complex sorting rule where cards with a black border are sorted by color and those without are sorted by shape (12 trials), and finally a rule switch to the opposite meaning of the border (12 trials). Participants advance to the more challenging sorting rules only if they meet criteria, i.e. 5/6 cards are successfully sorted by shape. In the current study, each correct trial was given a point for a maximum score of 36.

#### Sustained Attention (ECAB)

The Early Childhood Attention Battery (ECAB) is adapted from the Test of Everyday Attention for Children (TEACh) and validated for children aged 3-6 years (Breckenridge et al., 2013). Here we analyze data collected during the sustained attention subtest which showed children a series of pictures and asked them to identify each time an animal was presented (200 ms presentations with an inter-stimulus interval [ISI] of 1800 ms). Children are scored for each correct detection with a point deducted for each false positive, with a maximum score of 30.

#### Selective Attention (ECAB)

For this study we used the Apple Search subtest, which asks children to find a set of 18 red apples in a visual array that included white apple and red strawberry lures. Two sheets were generated, an ‘A’ version given at the baseline visit and a ‘B’ version at the follow up. This test is scored as the number of red apples correctly found in 60 seconds, with no points deducted for misidentifications.

#### Short-term and Working Memory

The Thorell Forwards and Backwards span tests (Thorell & Wåhlstedt, 2006) are adapted from Digit Span subsets off the Wechsler Intelligence Scale for children – 3^rd^ edition (WISC-3). The Thorell span tasks are especially suitable for young children as words are used as stimuli instead of digits (Thorell & Wåhlstedt, 2006). The words are simple nouns that most children know well (e.g., cat, tree, rabbit, and clown) and the series of words is to be produced first in a forward (short-term memory) and then in a backwards (working memory) order to that presented by the experimenter. Children were presented with two trials at each level and the test ended if children were incorrect on both trials at a given level. One point is awarded for each correct trial (i.e., producing all words in the correct order) and the sum of points was used as a measure of working memory (Thorell & Wåhlstedt, 2006). The maximum possible score was 12 (2 tries at 2-7 words) for the forward and 8 (2 tries at 2-5 words) for the backward tests.

### Statistical Analyses of demographic and outcome data

Statistical analyses were performed in RStudio (Team, 2023). Demographics were assessed using descriptive statistics to characterize the sample and identify any differences between children assigned to the DI or EG groups. We considered chronological age, gestational age at birth, birth weight, sex, race/ethnicity, as well as parent schooling as an indicator of socioeconomic status. Group comparisons for demographics were assessed using two-sample t-tests for continuous variables such as age, and chi square analyses for categorical variables. Attention and working memory scores were compared pre-/post-intervention using repeated measures ANCOVAs with visit and group as effects of interest and age, sex and gestational age as covariates. Forced choice (yes/no) responses to exit interview questions were compared between groups using chi-squared tests, and an independent samples t-test was used to compare a summarized measure.

### Qualitative description of open-text exit interview responses

Exit interviews included open-text questions that were transcribed. Illustrative quotes were identified and reported to provide parent perspectives on acceptability, barriers and facilitators.

## Results

### Participant Demographics

Demographic characteristics for both groups are presented in Table 1. Participants in the EG group were younger and had a lower mean FSIQ than the DI groups.

**Table 1:** Demographics for DI and EG groups. Groups were compared using two-sample t-tests and chi -square tests with difference at p<0.05 indicated with *. + Birthweight data was missing for two DI participants.

| Demographic Variables |  | DI<br>N=10 | EG<br>N=12 | *p* |
| --- | --- | --- | --- | --- |
| Age in years at baseline: M (SD) |  | 4.99 (0.48) | 4.58<br>(0.39) | 0.039 * |
| Sex: M/F |  | 5/5 | 5/7 | 1.000 |
| Gestational age at birth: M (SD) |  | 29.84 (3.11) | 32.13<br>(2.82) | 0.086 |
| Birth weight+: M (SD) |  | 1441.63 (586.14) | 1717.25<br>(687.70) | 0.365 |
| FSIQ: M (SD) |  | 111.50 (19.02) | 95.75<br>(11.38) | 0.026 * |
| Conner's Early Childhood Total T-Score |  | 58.9 (12.6) | 65.6<br>(14.1) | 0.23 |
| Race/ethnicity | White/European/Caucasian | 5 | 8 | 0.206 |
|  | Other | 5 | 2 |  |
|  | White/European/Caucasian+Other | 0 | 2 |  |
| Primary language spoken at home | English | 9 | 11 | 0.714 |
|  | Other | 1 | 0 |  |
|  | English/Other | 0 | 1 |  |
| Highest parental education completed | Did not complete High School | 0 | 1 | 0.561 |
|  | High School | 0 | 1 |  |
|  | Technical/Trade | 0 | 2 |  |
|  | Bachelor's Degree | 7 | 5 |  |
|  | **Master's Degree** | 3 | 3 |  |

### Feasibility and fidelity

A review of the per-session tracking sheets indicated that games were played close to the recommendation with some variation across families: recommended was 2–3 games per session, 3 times a week, actual was 1-5 games per session, 0-4 times per week depending on the family and week. Overall, this suggests feasibility for families to schedule in time for this intervention. We noted that some participants played additional DI intervention games despite only three out of the five games being recommended for this study. When comparing time spent on intervention, we found that participants in the EG group played significantly more sessions and an overall longer amount of time relative to the DI group (Table 2).

**Table 2:**
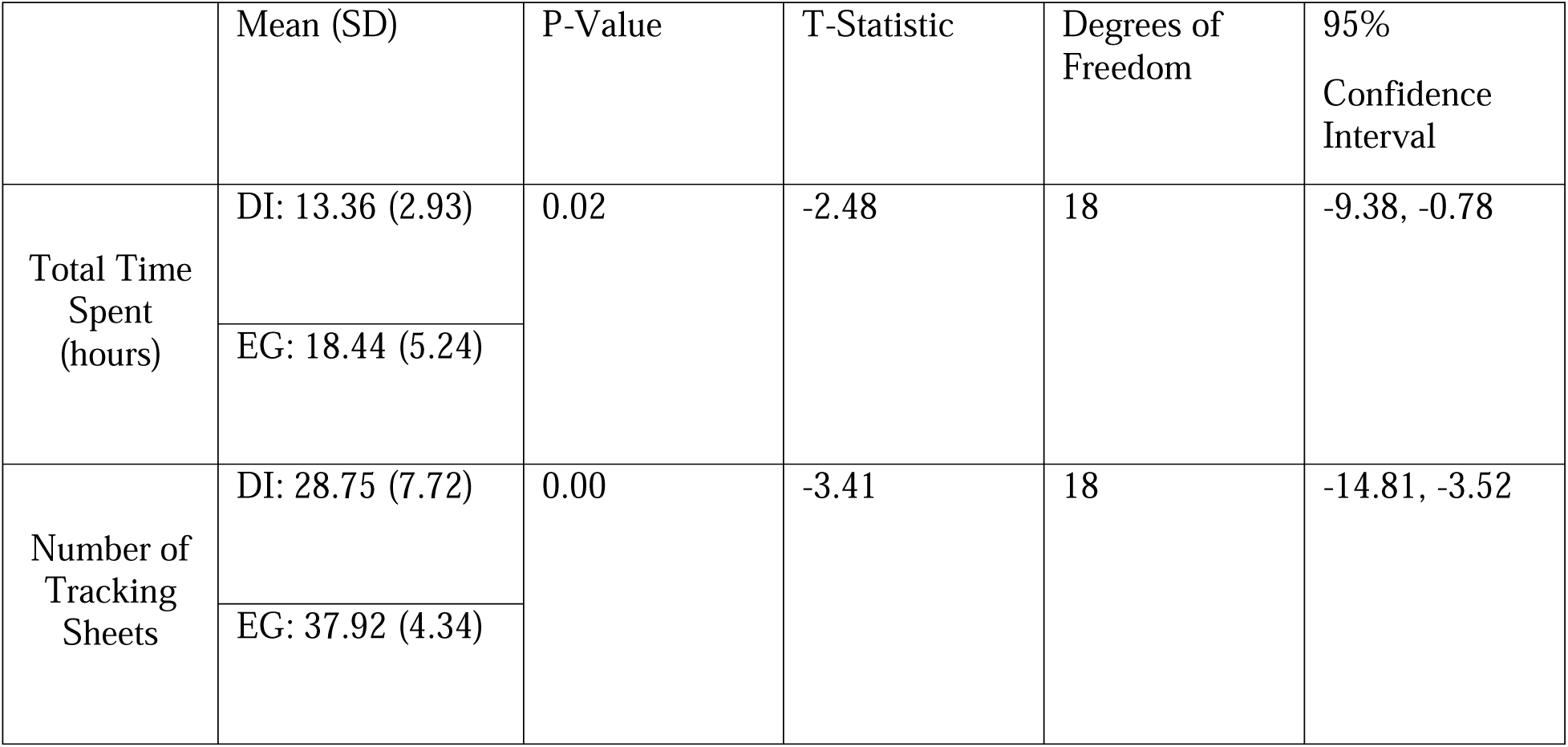
Intervention Statistics for DI vs. EG; The total time spent (hours) on DI or EG and the number of tracking sheets submitted by families from each group are shown here, along with group comparisons. Tracking forms were missing for two DI participants, for N=8 reporting in DI and N=12 in EG groups.

|  | Mean (SD) | P-Value | T-Statistic | Degrees of Freedom | 95% Confidence Interval |
| --- | --- | --- | --- | --- | --- |
| Total Time Spent (hours) | DI: 13.36 (2.93) | 0.02 | -2.48 | 18 | -9.38, -0.78 |
|  | EG: 18.44 (5.24) |  |  |  |  |
| Number of Tracking Sheets | DI: 28.75 (7.72) | 0.00 | -3.41 | 18 | -14.81, -3.52 |
|  | EG: 37.92 (4.34) |  |  |  |  |

Video calls with parents showed that all parents in the DI group were using some metacognitive strategies during intervention sessions. Tracking sheets from parents in the DI group further showed that metacognitive strategies were delivered alongside DI as instructed. As noted in Table 3, use was varied across families but all reported some use of metacognitive strategies. The frequency of strategy use is shown in Figure 1, with the most commonly used strategies shown to be self-awareness/monitoring, self-control and good listening.

**Figure 1.**
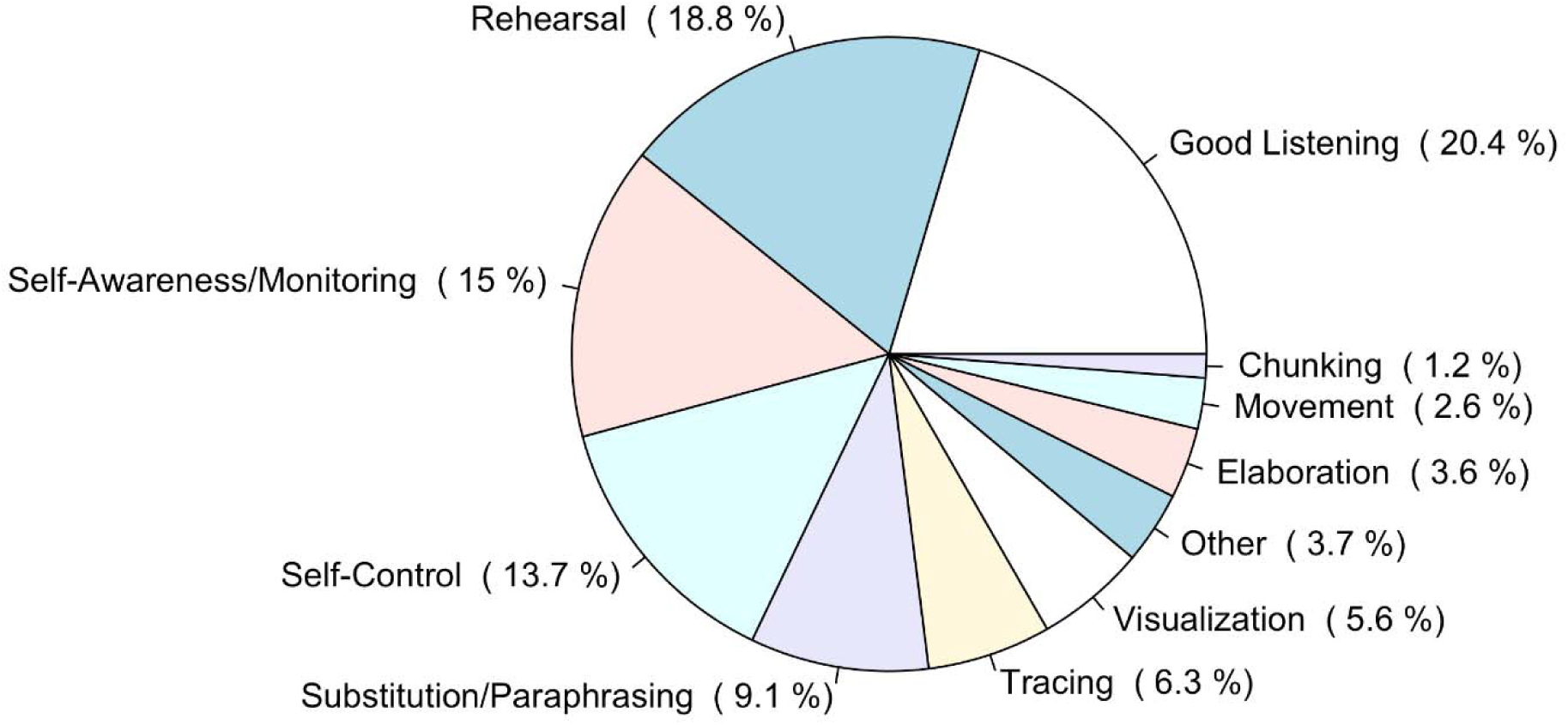
Frequency of Metacognitive Strategy Use. Metacognitive strategies included in ‘other’ were calming strategies, pattern recognition, pointing, talking aloud to think, and modelling.

**Table 3:** Total and average number of cognitive strategies reported by DI participants. Tracking forms were missing for two DI participants.

|  | **Average** | **SD** | **Min.** | **Max** |
| --- | --- | --- | --- | --- |
| Total Number of Cognitive Strategies Mentioned by DI Participants | 140.13 | 146.40 | 26 | 376 |
| Average Number of Cognitive Strategies Applied per Session by DI Participants | 4.26 | 3.74 | 0.93 | 10.16 |

### Intervention outcomes at 3-month follow-up

Selective attention, sustained attention, short-term and working memory assessments showed a significant effect of Visit, suggesting a practice or familiarity effect on change in scores over the three-month period. We also found an overall effect of Group for sustained attention and working memory, perhaps suggesting differences in baseline functioning that were maintained. While none of the scores showed a significant Group x Visit interaction effect, the DCCS executive function task showed a trend-level effect where children in the DI group showed greater gains (p=0.076). There was also a trend in the Thorell Forward short-term memory task where the EG group showed a greater positive change over time than the DI group (p=0.07; Table 4).

**Table 4.**
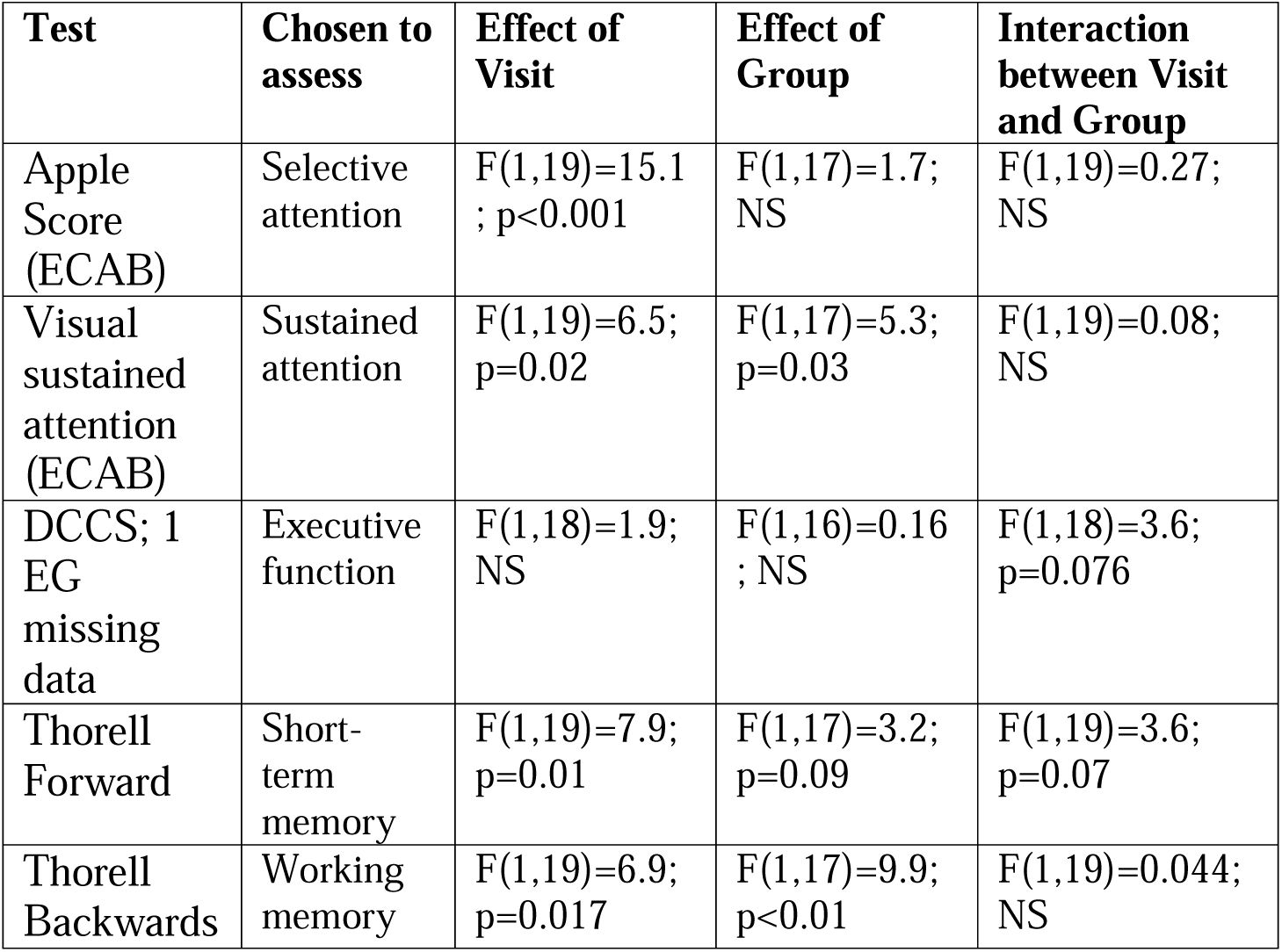
Statistical analyses of changes in cognitive scores across visits and interaction with intervention group. P-values are uncorrected for multiple comparisons.

| **Test** | **Chosen to assess** | **Effect of Visit** | **Effect of Group** | **Interaction between Visit and Group** |
| --- | --- | --- | --- | --- |
| Apple Score (ECAB) | Selective attention | F(1,19)=15.1 ; p<0.001 | F(1,17)=1.7; NS | F(1,19)=0.27; NS |
| Visual sustained attention (ECAB) | Sustained attention | F(1,19)=6.5; p=0.02 | F(1,17)=5.3; p=0.03 | F(1,19)=0.08; NS |
| DCCS; 1 EG missing data | Executive function | F(1,18)=1.9; NS | F(1,16)=0.16 ; NS | F(1,18)=3.6; p=0.076 |
| Thorell Forward | Short-term memory | F(1,19)=7.9; p=0.01 | F(1,17)=3.2; p=0.09 | F(1,19)=3.6; p=0.07 |
| Thorell Backwards | Working memory | F(1,19)=6.9; p=0.017 | F(1,17)=9.9; p<0.01 | F(1,19)=0.044; NS |

### Exit interview forced-choice questions

When asked yes/no questions about observed changes in behavior, parents in both groups noted some changes through the intervention. While reported changes tended to be more frequent in the DI group, differences generally did not reach statistical significance, with the exception that significantly more families in the DI group endorsed seeing their children apply strategies to help them focus (Table 5).

**Table 5:** Exit Interview Responses. One DI and 1 EG participant did not complete an exit interview.

| **Question:** |  | **Number of DI Parents Observing a Change in Behaviour** | **Number of EG Parents Observing a Change in Behaviour** | **P (chi-squared)** |
| --- | --- | --- | --- | --- |
| “Did you notice any changes (in your child) in these particular areas:” | Attention/Focus | 6/9 (66.67%) | 4/11 (36.36%) | 0.37 |
|  | Regulation of Emotions | 5/9 (55.56%) | 3/11 (27.27%) | 0.36 |
|  | Regulation of Behaviour | 4/9 (44.44%) | 3/11 (27.27%) | 0.64 |
|  | Social Skills | 1/9 (11.11%) | 3/11 (27.27%) | 0.59 |
|  | Initiation | 4/9 (44.44%) | 2/11 (18.18%) | 0.34 |
|  | Task Completion | 6/9 (66.67%) | 5/11 (45.45%) | 0.41 |
|  | Communication | 4/9 (44.44%) | 6/11 (54.55%) | 1.00 |
|  | Flexibility | 3/9 (33.33%) | 2/11 (18.18%) | 0.62 |
|  | Efficiency/Productivity | 3/9 (33.33%) | 2/11 (18.18%) | 0.62 |
|  | Impulsivity | 2/9 (22.22%) | 2/11 (18.18%) | 1.00 |
|  | Following Directions | 6/9 (66.67%) | 3/11 (27.27%) | 0.17 |
|  | Remembering | 6/9 (66.67%) | 6/11 (54.55%) | 0.67 |
| Average Total Score |  | 5.56 | 3.73 | 0.31 (t-test) |
| “Have you noticed your child applying any specific strategies to help him or her focus or remember since playing the games?” |  | 6/9 (66.67%) | 1/10 (10%); 2 EG did not respond | 0.02* |
| “Have you noticed your child applying specific metacognitive strategies when thinking/learning?” |  | 5/9 (55.56%) | 3/9 (33.33%); 3 EG did not respond | 0.64 |

### Exit interview open-text questions

Open-text responses on exit interviews were used to inform acceptability and far transfer benefits of metacognitive strategy learning from the DI intervention. Some parents reported changes in behavior following DI. Specifically: [my child was] “Able to follow directions better, the ability to finish more tasks and improved concentration.” and “I noticed her using the rehearsal strategy to ‘think aloud’ much more. She also created stories in her drawings a bit more than she did prior to the study…”. Additional beneficial impacts of metacognitive strategies that transferred to everyday behaviors were: “Repeating verbally helped [my child] remember multi-step tasks.” “[My child] is getting better at managing [] emotions, & [] overall patience has improved (waiting, following through).” “[My child] used elaboration aloud to demonstrate [] understanding of TV shows, read stories.”, “I’ve seen how ‘chunking’ long strings of information into smaller pieces helps [my child] remember, as well as focusing on the main aspects of items, ideas.”, “[My child] communicates aloud much better than [] prior to the games [and] seems more aware of noticing how/when to adapt [their] way of communicating & is a more flexible learner”.

In terms of acceptability, parents reported that children enjoyed the games. A facilitator is the rewards that are included in DI, specifically children enjoyed collecting coins and picking out prizes. Parent-child interactions were also noted as positive: “The ‘teaching [] how to think a way through trouble areas’ was a great takeaway. [My child] also gained, I think, some insight or increased awareness of how [] frustration ‘gets in the way’ of learning at times. [My child] learned some useful coping skills (breathing, changing gears).”

In terms of barriers to success, some parents reported experiencing some technical difficulties that will be remedied in future iterations. Among the 8 DI families who withdrew, two completed a questionnaire providing some context. From these families we heard that barriers included finding time to schedule the intervention and around excessive screen time for their child.

## Discussion

The present pilot study tested feasibility and preliminary efficacy of a parent-delivered combined process-specific and compensatory tablet game-based intervention for attention and working memory in early school-aged children who were born preterm. Key findings were: 1) parents can be effective interventionists and can successfully support their children in applying metacognitive strategies, 2) greater beneficial impacts on child behavior were found in the DI relative to the educational games control group, 3) we did not find a significant benefit of the DI intervention on performance-based measures of attention and working memory, and 4) there was greater attrition and less time on intervention in the DI relative to the EG control group, suggesting that ongoing refinement of the program may help to enhance feasibility in future. Together this work advances our understanding of opportunities for improving school-readiness in children who were born preterm.

The DI/CQ programs were initially developed to be administered by trained professionals such rehabilitation experts or educational assistants (Kerns et al., 2017; Kerns et al., 2010; Macoun et al., 2022; Macoun et al., 2021). This approach maximizes fidelity but limits the potential for the intervention to be offered to children who do not have sufficient resources at their schools or in their community. Parent-delivered is attractive in order to broaden the reach of the program and also, parent training can have downstream benefits for child development and parenting efficacy/skills (Kaminski et al., 2008). For the present study, we developed a parent training program that included: a slide presentation delivered in-person, regular check-ins with the study team and a video-call based fidelity check during which a research assistant observed delivery of the intervention and provided feedback. Based on the fidelity observation and tracking sheets, the DI intervention was largely delivered as intended in terms of frequency, duration and supporting metacognitive strategy use. Facilitators included compelling bonus games and barriers related to challenge with scheduling and screen time.

Despite overall success in implementation, we noted that parents in the DI group engaged with the intervention less frequently and spent less total time on the intervention than the EG control group. This perhaps reflects that DI is designed to be repetitive and challenging, whereas the games chosen in the EG group were not similarly repetitive or progressive. In addition, parents were learning new skills through delivering an intervention approach not typically offered in home settings. While this improves accessibility to cognitive interventions that have been traditionally difficult to access, adaptations to improve engagement and motivation will be critical. The success of the intervention in the control group can offer some insights for future intervention programs. Parents in the EG group generally reported that their children enjoyed the games and were happy to play. Parents in the DI group reported that the games were enjoyable but challenging and that further development of the bonus games would be helpful. The features of the educational games that make them enjoyable can be more deeply integrated into programs like DI going forward and the bonus games in DI could be expanded. Perhaps access to educational games could also be leveraged as a reward to motivate playing the more challenging DI games. Additional tools and support for parents to help their children to persist at challenging tasks and manage frustration, could be offered through weekly parent support groups during active intervention.

Given the potential for practice and placebo effects, we included a control group that included playing tablet games under parental supervision but did not include the specific elements of the DI program: hierarchical games that challenge attention and working memory skills and training in metacognitive skills to scaffold learning. We found that on several attention and working memory outcome measures, there were increases in scores across the entire sample that were not higher in the DI group. This highlights the importance of potential practice effects. On the other hand, we acknowledge that interactions require greater statistical power relative to main effects, and we may be simply underpowered to detect intervention-specific changes with the sample recruited here. Anecdotally, parents in the DI group reported behavioural changes in their children that were not seen in the educational games group including far-transfer of metacognitive strategies to settings outside of the game.

Our results add to a growing but mixed literature on cognitive training interventions for preterm and neurodevelopmental populations. Early pilot work demonstrated that computerized executive function training is feasible and can yield improvements in visuospatial working memory among very preterm children, although these findings were based on a small sample and prevent drawing firm conclusions about the efficacy of the training on said changes (C. S. H. Aarnoudse-Moens et al., 2018). However, subsequent larger studies have reported inconsistent effects. For example, van Houdt and colleagues (2020) found no significant improvements in attention, executive function, or academic outcomes following six weeks of game-formatted executive function training (i.e., BrainGame Brian) in very preterm children (van Houdt et al., 2021). Similarly, Anderson et al. (2018) reported that Cogmed Working Memory Training did not improve long-term academic functioning or attentional skills in extremely preterm children at 24-month follow-up (Anderson et al., 2018). These findings highlight the challenges of achieving robust and generalizable effects in this high-risk population, particularly when hybrid approaches are not. By contrast, meta-analytic evidence suggests that Cogmed can reduce inattentive behaviors in daily life across child and adult populations, including those with neurodevelopmental disorders [(Spencer-Smith & Klingberg, 2015). While these benefits may not consistently extend to broader academic or behavioral outcomes, they suggest that improvements in cognitive processes are possible with well-designed training protocols.

Evidence from de Vries et al. (2015), who tested executive function games in children with autism, highlights the challenges of keeping participants engaged. Although their study found some signs of improvement in working memory, high dropout rates and minimal differences from the control group limited the overall impact (de Vries et al., 2015). These issues reflect common difficulties in digital interventions, where gains on training tasks do not always extend to everyday functioning. In contrast, our DI approach included both strategy coaching and active parent involvement, which may help children apply the skills they practiced in the games to their daily lives. This emphasis on metacognitive support and generalization could explain why parents in our study observed benefits beyond the training itself.

Strengths of this study included a control group and validity checks, thorough parent training and fidelity checks, and use of a hybrid intervention in keeping with best-practice recommendations for cognitive training (Cicerone et al., 2011; Kennedy et al., 2008; Shipstead et al., 2012; Sohlberg et al., 2003; von Bastian & Oberauer, 2014). There are also several limitations to note. The small sample size is useful for demonstrating feasibility and tolerability of the program but is likely underpowered to detect meaningful cognitive changes. Other limitations include a restricted age range and lack of long-term follow-up to investigate maintenance of gains over time and longer-term impacts on academic function.

In summary, we present here a study that demonstrates feasibility of a parent-delivered tablet game-based intervention in children who were born prematurely. While we generally found that the intervention was feasible in this format, and parents reported improvements in behavior, we also found that there is room for improvement in the game delivery. We further did not find a specific impact of the DI games on change in attention and working memory measures in this small sample. Children born preterm are vulnerable to challenges with school readiness. Intervention studies, such as this, are important for advancing our ability to support children at greatest risk.

